# Co-evolution between codon usage and Protein-Protein Interaction Networks in Bacterial genomes

**DOI:** 10.1101/2020.03.30.016956

**Authors:** Maddalena Dilucca, Giulio Cimini, Sergio Forcelloni, Andrea Giansanti

**Affiliations:** Dipartimento di Fisica, Sapienza University of Rome, Rome, Italy; Dipartimento di Fisica, Tor Vergata University of Rome, Italy; Istituto dei Sistemi Complessi (CNR) UoS; INFN Roma1 unit, Rome, Italy

**Keywords:** codon usage bias, bacteria, protein-protein interaction networks, interactomes

## Abstract

In this work, we study the correlation between codon usage and the network features of the PPI in bacteria genomes. We want to extend the information by Dilucca et al. (2015) about E.Coli’s genome for a set of other 34 bacteria. We use PCA techniques in the space of codon bias indices (compAI, compAI_w, tAI, NC) and GC content to show that genes with similar patterns of codon usage feature have a significantly higher probability that their encoded proteins interact within the PPI. And vice-versa, we show that interacting in the PPI have a coherent codon usage. This work could allow for future investigations into the possible effects that codon bias signal can have on the topology of protein interaction network and, as such, to improve existing bioinformatics methods for predicting protein interactions.

## 1. Introduction

In this work, we investigate the relationship between codon bias of coding sequences, at the genetic level [1] and connectivity patterns of protein-protein interaction networks (PPI) [2].

As it is well-known, the genetic code is degenerate: synonymous codons at the genetic level encode for the same amino acid in the translated protein. Moreover, although synonymous codons are indistinguishable in the primary structure of a protein, they are not used randomly, but with different frequencies which may vary across different species, in various parts of one specific genome, and even within different regions of the same gene This phenomenon, known as Codon Usage Bias (CUB), is well-established in the literature (see, for example, [3-5]). Nevertheless, while remarkable observations have emerged, a general understanding of the biology of CUB still lacks, because of its complex phenomenology [7]. Although CUB does not alter the amino acid sequences of proteins, it is involved in many important cellular processes, including differential gene expression [7], translation efficiency and accuracy [8,9], dynamics of the ribosome, and co-translational folding of the proteins [10,6].

With the present study, we intend to share basic observations of sufficient generality on the co-evolution of CUB and the degree connectivity of bacterial interactomes. The systematic analysis of PPIs is crucial to understand the patterns of chemical reactions within cells, and the role played by proteins in regulative processes. Moreover, on the applicative side, the comparison of the interactomes from different species is fundamental to understand disease-related processes that engage more than one species, such as host-pathogen relationships, to identify clinically relevant host-pathogen PPI, and consequently to develop future therapeutic applications [1]. In previous work, Dilucca et al. showed that translational selection in E. Coli (taken as a case of study) systematically favors optimal codons in proteins that have a large number of interactors and belong to the most representative communities in the PPI [3]. In the current analysis, we extend those observations to a set of unrelated bacterial species by focusing on the connection between the codon usage of the genes and the role of their corresponding proteins in the protein interaction network. It is well known that codon usage bias is strongly correlated with expression level in many organisms, and it is well structured along the genome, with near neighbor genes having similar usage frequencies of synonymous codons [13]. Accordingly, and considering that gene expression level [14] and proximity between the positions of the genes in the genome are powerful predictors of protein-protein interaction, we analyze how the closeness in codon usage bias of the genes (as measured by the normalized scalar product of RSCU vectors) is reflected into the capacity of the corresponding proteins to make physical contact in the cell. Given the above considerations, we would expect a more similar codon usage bias between interacting proteins than non-interacting ones. We find out that this statement is not valid for all bacteria considered in this analysis, especially if we measure CUB without taking into account the information about tRNA levels. This means that the distance in codon usage bias seems unable to distinguish between proteins that make contacts or not in the protein interaction network. To further investigate this issue, we performed a Principal Component Analysis (PCA) in the space of the codon bias indices (NC, tAI, compAI, GC content), thus using compAI and tAI to add the information about the tRNA pool of the organism. In all bacteria, we observed a clear localization of the most representative communities of the protein interaction networks in the space of the first two PCA components, underlining a more pronounced similarity between proteins belonging to the same community. This observation could be explained considering the fundamental requirement of interacting proteins (especially those belonging to the same community) to be present in the cell according to precise quantities at a given time to form the protein complexes that are necessary upon the ongoing cellular programs. Finally, we show that the distance between a pair of genes in the plane of the first two PCA component is a statistically robust predictor of the likelihood that their corresponding proteins do interact (physically or functionally). In agreement with our previous results, the signal is evident when codon usage frequencies of interacting proteins are far from being random.

To summarize, in line with what observed in the first work [3], our main result indicates that the functional structuring of the protein interaction network has interfered with the peculiar codon choice of the genes over evolutionary time. These results point out that codon bias evolution should be a relevant parameter in the prediction of unknown protein-protein interactions from only genomic information. Last but not least, we claim that using the codon usage bias as an additional level of information in the study of protein interaction networks could be useful to provide alternative treatments in the light of growing resistance to antibiotics and the propagation of numerous infectious agents.

## 2. Materials and Methods

### 2.1 Data sources

In this work, we select a set of 34 bacterial genomes with different behavior, environment and taxonomy (see Table 1 for details). Each bacteria represents a specific clade in the phylogenetic three by Plata et al. [14]. Nucleotide sequences were downloaded from the FTP server of the National Center for Biotechnology Information (ftp://ftp.ncbi.nlm.nih.gov/genomes/archive/old_genbank/Bacteria/) [2].

**Table 1.**
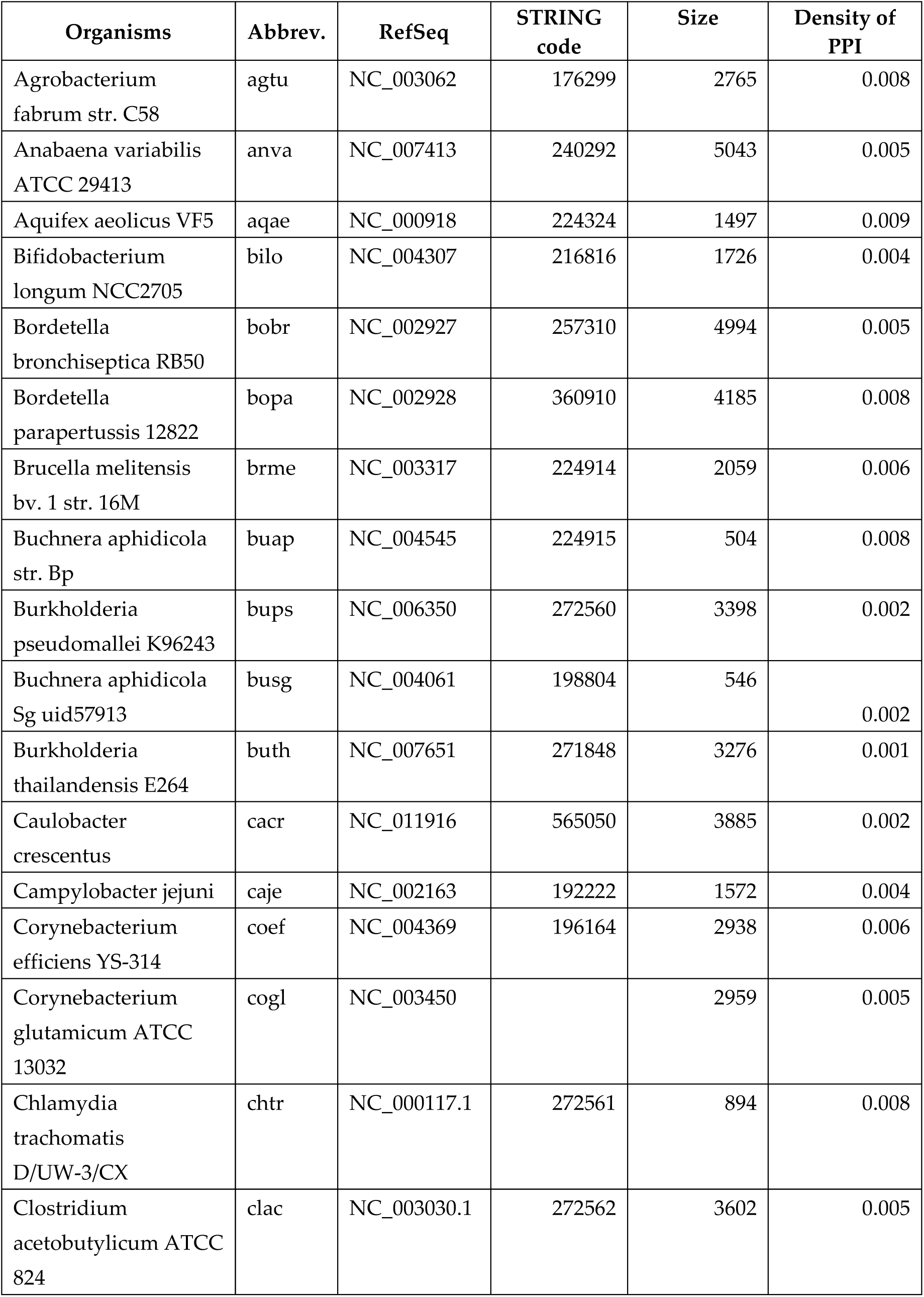
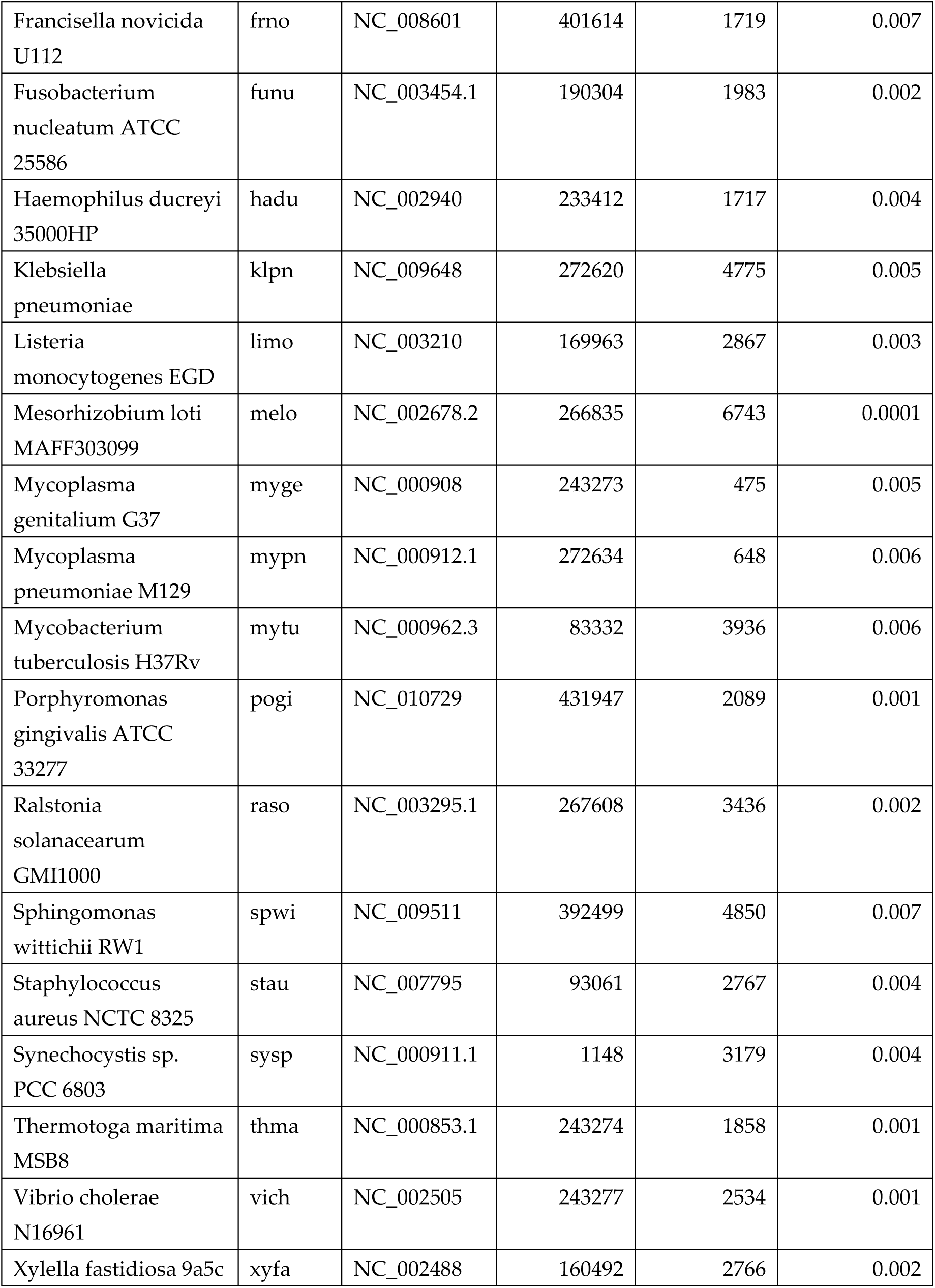
Summary of the 34 bacterial dataset considered in this work. In table, we report the organism name, abbreviation, RefSeq, STRING code, Size of genome (number of genes), and the density of the protein interaction network defined as ratio between the number of links in the real interactome and the maximum number of possible links (n(n-1)/2, where n is the number of nodes).

### 2.2 Codon Usage Bias Measures

In this work, we used four measures to quantify the extent of CUB: 1) the tRNA Adaptation index (tAI) [5], which is based on the assumptions that tRNA availability is the driving force for translational selection; 2) CompAI and CompAI_w [3], which refine tAI using the competition between cognate and near-cognate tRNA; 3) the Effective Number of Codons (NC) [18], which is a statistical measure of the number of codons used in the sequence; and 4) the GC content, namely the percentage of guanine and cytosine in the RNA molecules. Moreover, we use the Relative Synonymous Codon usage (RSCU) to have a finer description of the non-uniform usage of synonymous codons in each coding sequence in the form of a 61-component vector.

### 2.3 Protein-Protein Interaction Network

PPI networks of the 34 bacterial genomes were retrieved from the STRING database (Known and Predicted Protein-Protein Interactions) [11]. Community detection in PIN was performed by using the Molecular Complex Detection (MCODE) software. In this study, we evaluate communities based on the k-core concept, i.e., the largest subgraph within which each node has at least K connections. In line with our previous study [3], we consider only the first eight communities with a degree k > 10.

### 2.4 Analyses of the RSCU vectors of genes encoding for interacting proteins in the PPI

The main advantage of RSCU is to allow quantifying the similarity between two genes in their usage of synonymous codons using the classical operations between vectors. This feature makes it particularly suitable for the study of how the similarity between two genes in terms of codon usage bias is reflected in the ability of their encoded proteins to make contact in the protein interaction network. Specifically, a vector of RSCU for each gene of a genome was constructed with a homemade Python script. We computed the normalized distribution of the scalar products between RSCU vectors for genes whose encoded proteins are in contact in the protein interaction network. We then compared this distribution with that obtained by considering a random model of the protein interaction network, in which all connections are randomly shuffled, but the number of connections of each protein is kept constant. For each pairwise comparison, we used the Mann-Whitney test for assessing if distributions are different (p-value < 0.001).

### 2.5 Principal Component Analysis

Principal Component Analysis (PCA) is a multivariate statistical method that transforms a set of possibly correlated variables into a set of linearly uncorrelated ones (called principal components, spanning a space of lower dimensionality) [6].

We use PCA over the space of the five codon bias indices described above. Thus, each gene is represented as a 5-dimensional vector with coordinates (compAI, compAI_w, tAI, NC, GC).

The first two principal components (*PC*_1_*and PC*_2_) turn out to represent as much as 65% of the total variance of codon bias over the genomes. Thus, we focus on the plane defined by these two vectors. We calculate this plane for each bacterial genome (see Figure 2 in Supplementary Materials), reporting the associated variances, eigenvalues and eigenvectors in Supplementary Materials. We then project the first eight communities extrapolated with MCODE in the *PC*_1_ − *PC*_2_ plane, and we calculate their centroids (with error bars defined by standard deviations) to check if these communities are well separated on this plane.

### 2.6 Z-score analysis

We define *d* as the Euclidean distance between two genes on the *PC*_1_ − *PC*_2_ plane, and Pr(*link*|*d*) as the fraction of gene pairs, among those localized within a distance *d*, whose encoded proteins are connected in the PPI network. We then compare this conditional probability estimated on the real interactomes with Pr(< *link* >_*Ω*_ |*d*), namely the same probability estimated on the configuration model (CM) of the networks (a degree-conserving rewiring of the network links) [22].

The CM is used as the null hypothesis that no relation exists between the codon usage of two genes and a physical interactions between the encoded proteins. The significance of Pr(*link*|*d*) with respect to the null hypothesis is quantified through the Z-score values:

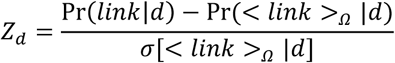

We computed the function of Z-score as a function of the gene distance *d* for each genome.

## 3. Results

### 3.1 Interacting proteins do not share a common codon usage statistics

We analyze how the closeness in codon usage bias of two genes (as measured by the normalized scalar product of their RSCU vectors) is reflected in the capacity of their proteins to make physical contact in the cell. According to what stated in the Introduction section, we expect a more similar codon usage bias between interacting proteins than non-interacting ones. Noteworthy, we find that this statement is not valid for all bacteria considered in this analysis if we measure CUB by not taking into account the information about tRNA levels. In other words, the pure codon statistics is not able to distinguish between proteins that are in contact or not in the protein interaction network. In Table 2, we report in boldface the 12 bacteria that pass the test (p-values < 0.001, Mann-Whitney test).

**Table 2.** P-values of the pairwise comparisons between the normalized distribution of scalar products of RSCU vectors for genes corresponding to interacting proteins in the PPI and distribution obtained by shuffling the PPI. For each bacterial dataset, we report the organism name, abbreviation, and p-value. The bacterial species for which the difference is statistically significant, according to the Mann-Whitney test (p-value < 0.001), are reported in boldface.

| Organisms | Abbr. | p-value |
| --- | --- | --- |
| **Agrobacterium tumefaciens** | agtu | 1.28*10<sup>-6</sup> |
| **Anabaena variabilis ATCC 29413** | anva | 1.02*10<sup>-4</sup> |
| Aquifex aeolicus VF5 | aqae | 0.20 |
| **Bifidobacterium longum NCC2705** | bilo | 0.15 |
| **Bordetella bronchiseptica RB50** | bobr | 0.27 |
| **Bordetella parapertussis 12822** | bopa | 1.06*10<sup>-3</sup> |
| **Brucella melitensis bv. 1 str. 16M** | brme | 1.56*10<sup>-7</sup> |
| **Buchnera aphidicola str. Bp** | buap | 0.13 |
| **Burkholderia pseudomallei K96243** | bups | 1.16*10<sup>-6</sup> |
| Buchnera aphidicola Sg uid57913 | busg | 0.035 |
| **Burkholderia thailandensis E264** | buth | 9.1*10<sup>-15</sup> |
| Caulobacter crescentus | cacr | 0.15 |
| Campylobacter jejuni | caje | 0.10 |
| **Corynebacterium efficiens**<br>YS-314 | coef | 0.27 |
| **Corynebacterium glutamicum**<br>ATCC 13032 | cogl | 6*10<sup>-3</sup> |
| Chlamydia trachomatis<br>D/UW-3/CX | chtr | 0.18 |
| **Clostridium acetobutylicum**<br>ATCC 824 | clac | 7*10<sup>-11</sup> |
| Francisella novicida U112 | frno | 0.66 |
| **Fusobacterium nucleatum**<br>ATCC 25586 | funu | 1.1*10<sup>-11</sup> |
| **Haemophilus ducreyi 35000HP** | hadu | 0.57 |
| **Klebsiella pneumoniae** | klpn | 0.89 |
| **Listeria monocytogenes EGD** | limo | 1.3*10<sup>-5</sup> |
| Mesorhizobium loti MAFF303099 | melo | 0.01 |
| **Mycoplasma genitalium G37** | myge | 5.8*10<sup>-5</sup> |
| **Mycoplasma pneumoniae M129** | mypn | 5.2*10<sup>-5</sup> |
| Mycobacterium tuberculosis<br>H37Rv | mytu | 1.6*10<sup>-3</sup> |
| **Porphyromonas gingivalis**<br>ATCC 33277 | pogi | 4.5*10<sup>-6</sup> |
| Pseudomonas aeruginosa<br>UCBPP-PA14 | psae | 0.016 |
| Ralstonia solanacearum GMI1000 | raso | 0.022 |
| Rickettsia prowazekii str. Madrid<br>E | ripr | 0.18 |
| **Sphingomonas wittichii RW1** | spwi | 8.9*10<sup>-9</sup> |
| Staphylococcus aureus NCTC<br>8325 | stau | 0.02 |
| **Synechocystis sp. PCC 6803** | sysp | 3*10<sup>-4</sup> |
| **Thermotoga maritima MSB8** | thma | 1.6*10<sup>-3</sup> |
| **Vibrio cholerae N16961** | vich | 2.1*10<sup>-8</sup> |
| **Xylella fastidiosa 9a5c** | xyfa | 5.1*10<sup>-4</sup> |

### 3.2 Principal Component Analysis over the space of the codon bias indices

We then perform PCA over the space of the five codon bias indices (CompAI, CompAI_w, tAI, NC and GC content) measured separately for each gene in each genome (see Figure 1 for an example). The other plots are shown in Supplementary Material. The two first principal components (*PC*_1_ and *PC*_2_) turn out to represent for as much as 65% of the total variance of codon bias over each genome (see plot of Figure 2 for example). The other plots are shown in Supplementary Material. Projection of the first two principal components on the individual codon bias indices (loadings) for each genome shows that none of the five indices predominantly contributes to the data variability (see plot Figure 2 for example). Thus, the placement of a gene in the *PC*_1_ − *PC*_2_ plane depends on a weighted contribution of all the indices. So, we calculate the centroids of the eight top MCODE communities in this plane. We show that the genes encoding for the respective proteins are well localized in this reduced space (see Figure 1). For each genome, the first community (composed overall by 97 % from genes class COG J, that are translational, ribosomal structure and biogenesis) is always separated and isolated from the others. For the other communities, it depends on the bacteria: some bacteria such as *raso, ripr* or *bups* have all communities well separated, whereas other bacteria such as *caje, chtr* or *pogi* do not have separated centroids (see Figure 2 of Supplementary Material). In other words, for some bacteria, if a set of proteins are physically and functionally connected in a module, then their corresponding genes should share common codon bias features.

**Figure 1.**
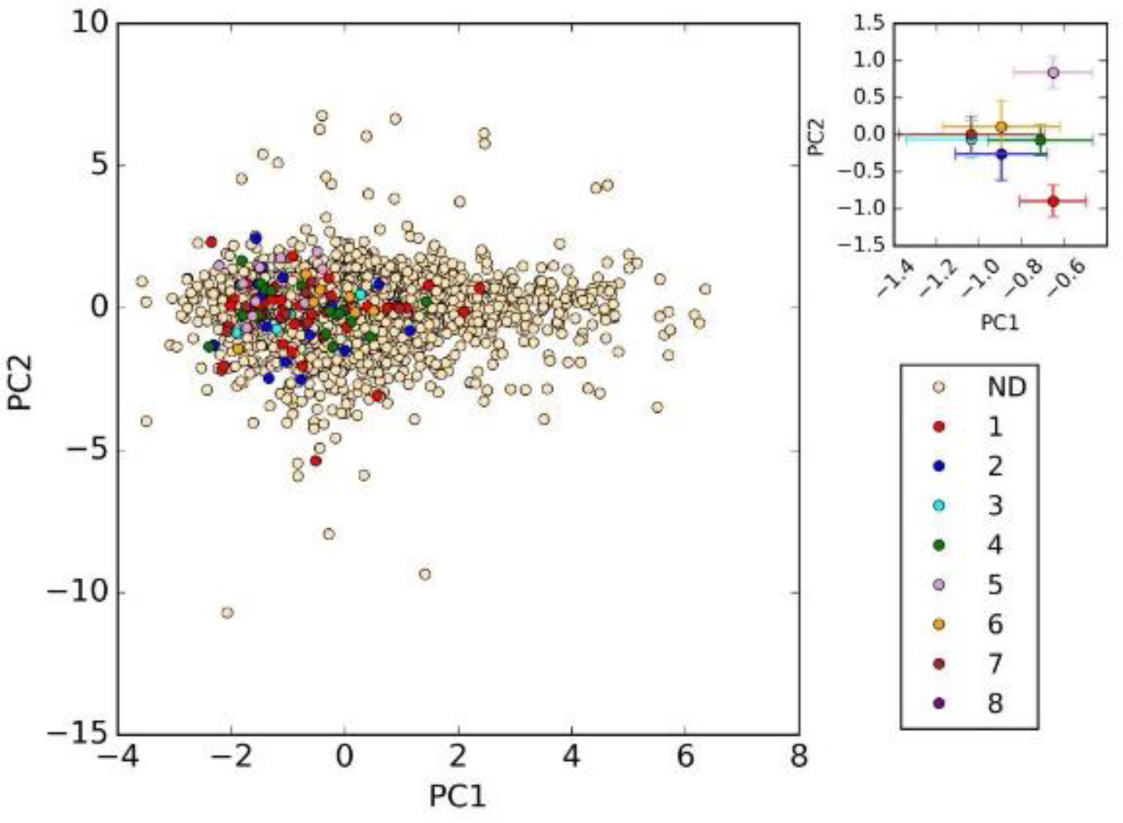
Example of plane *PC*_1_ − *PC*_2_. We show the example of agtu species. In the inset, there are centroids of first eight communities. We show that first community is always well separated from the others. The other bacteria are shown in Supplementary Material.

**Figure 2.**
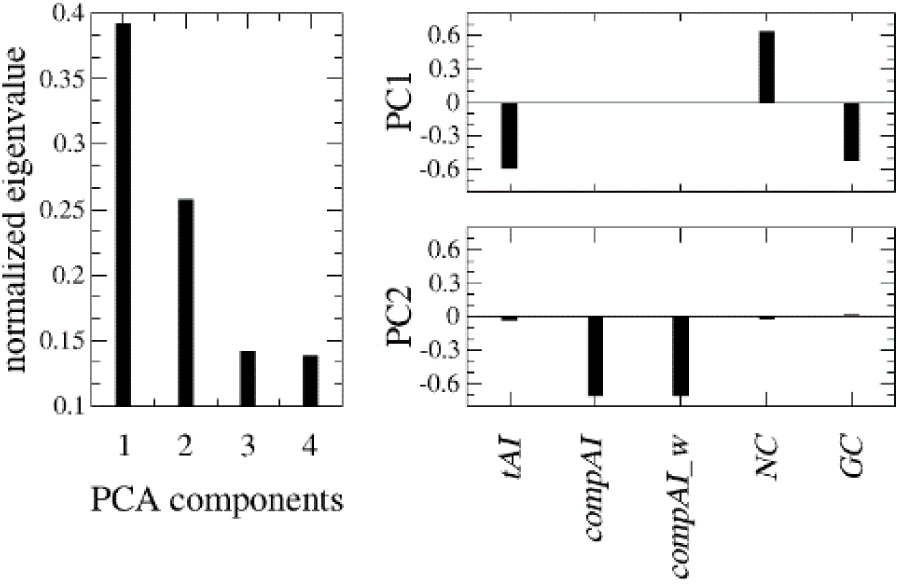
Example of eigenvalues of the correlation matrix. We show the example of agtu species. The first and second principal components (*PC*_1_ − *PC*_2_) turn out to represent as much as 68% of the total variance of codon bias. The other bacteria are shown in Supplementary Materials.

### 3.3 Z-score profiles: the closer is the codon usage of genes, the higher is the probability of protein interaction

Finally, for each bacterial genome, we calculate a conditional probability *Pr*(*link*|*d*) of a functional interaction between proteins, given that their relative genes fall within a distance *d* in the plane of the two principal components *PC*_1_ − *PC*_2_ [2]. We compare *Pr*(*link*|*d*) estimated on the real interactomes downloaded from STRING with Pr(< *link* >_*Ω*_ |*d*) estimated using the Configuration Model (CM) of the network, in which links are shuffled by preserving the interaction degree of each protein (see Materials and Methods for details).

To compare these two conditional probability, we study the Z-scores for *Pr*(*link*|*d*)) as a function of the Euclidean distance *d* between the codon usage bias of pair of genes in the PCA plane (see section. In Figure 3, we show the averaged Z-score profile over the set of bacteria considered in this analysis.

**Figure 3.**
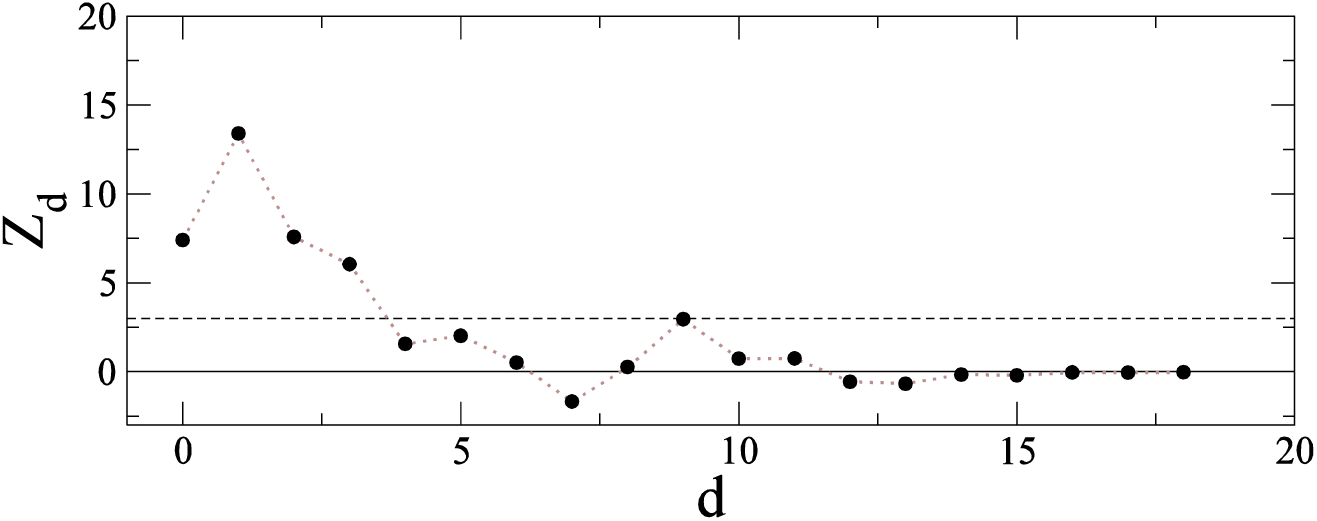
Z-scores of *Pr*(*link*|*d*) as a function of the Euclidean distance d between the codon usage bias of pair of genes (computed via PCA), averaged over the set of chosen bacteria. The horizontal dashed lines mark the significance interval of 3σ standard deviations. We show the example of *agtu* species.

Interestingly, for small distances (d <= 3), the probability of finding a connection between two proteins in the real interactome is much higher than in the CM (Z-score > 1.96). Conversely, for larger distances (d > 3) the real network and the CM become compatible (−1.96 < Z-score < 1.96).

Looking at the individual profiles in Supplementary Material, we note that this profile is common among different bacterial species. Thus, as a general rule, we conclude that sets of genes sharing similar codon usage patterns encode for proteins that are more likely to interact in the protein interaction network than expected by chance. Notably, for all the bacteria that pass the test for RSCU distributions in Table 2, the individual Z-score profiles are similar to Figure 3. This is also valid for some bacteria (i.e., *busg, cacr, caje, coef, hadu, klpn, raso*, and *stau*) that instead do not pass the test in Table 2. In contrast, the rest of the bacteria (i.e., *aqae, bilo, bobr, buap, chr, frno* and *melo*) are characterized by a different Z-score profile (See Supplementary Material), probably due to the low density of their PPI (See Table 1).

## 4. Discussion

In this work, we study how the coherence in codon usage among genes is reflected in the capacity of the encoded proteins to interact in the protein interaction network. For this purpose, we have extended our previous work taking E. Coli as a case of study [3] to other 34 bacterial genomes characterized by different taxonomy [14]. As a general rule, we found that the only codon statistics based on the frequencies of occurrence of synonymous codons are not able to distinguish between proteins that make contacts or not in the protein interaction network. Conversely, by inserting the information contained in the tRNA levels, as expressed by tAI, CompAI, and CompAI_w, we observed that highly connected proteins belonging to the same communities in the protein interaction network are encoded by genes that are coherent in their codon choices. Specifically, our results provide evidence that if two genes have similar codon usage patterns, then the corresponding proteins have a significant probability of being functionally connected and interacting. Consequently, this study provides new information based on the similarity in codon usage of genes that can be potentially integrated into existing computational prediction methods of protein-protein interaction.

