## Supplemental file pdf for "Co-evolution between codon usage and Protein-Protein Interaction Networks in Bacterial genomes"

### Supporting Information

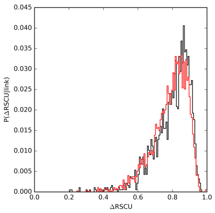

aqae

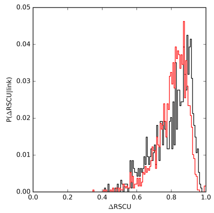

aqae

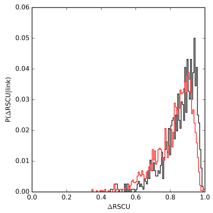

aqae

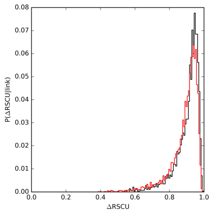

aqae

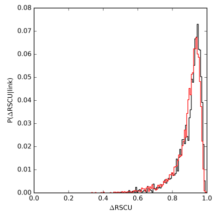

aqae

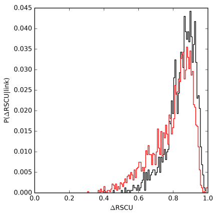

aqae

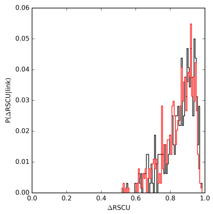

aqae

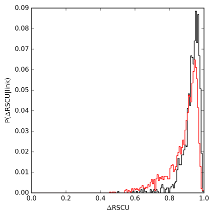

bups

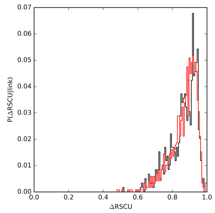

busg

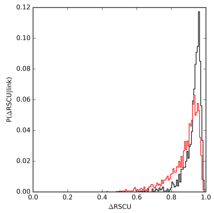

buth

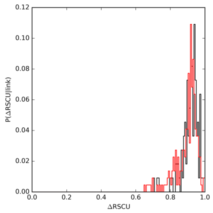

cacr

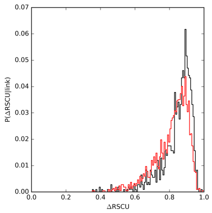

caje

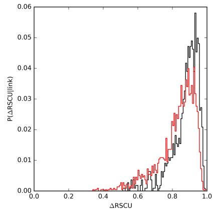

aqae

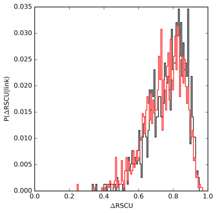

aqae

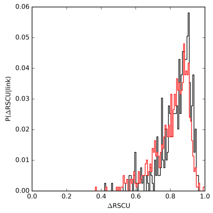

chtr

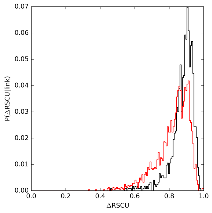

clac

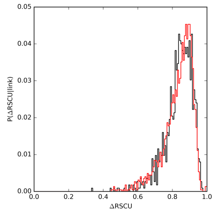

frno

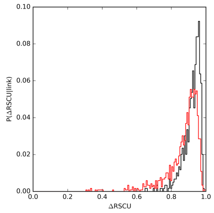

funu

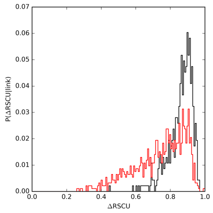

aqae

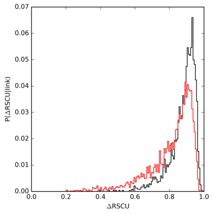

aqae

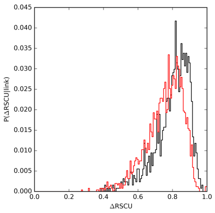

funu

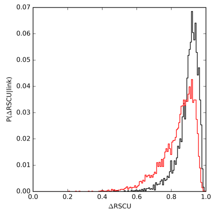

melo

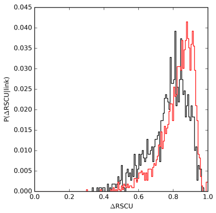

myge

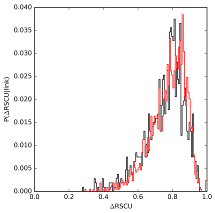

mypn



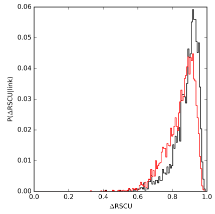

[illegible]

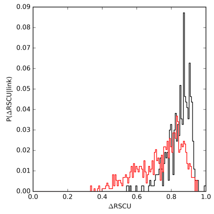

[illegible]

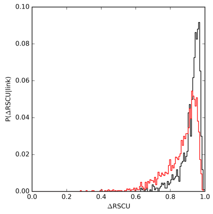

[illegible]

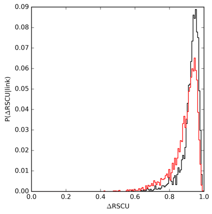

[illegible]

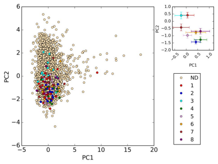

aqae

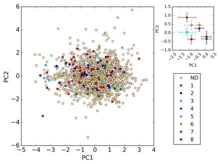

aqae

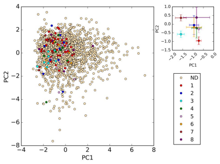

aqae

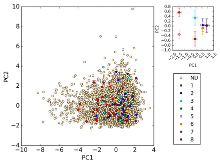

aqae

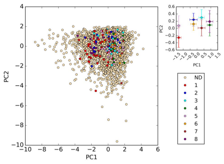

aqae

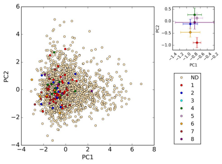

aqae

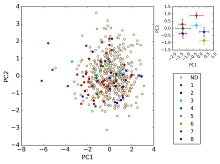

aqae

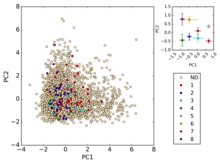

bups

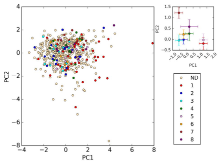

busg

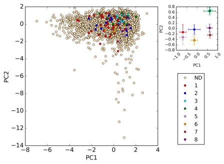

buth

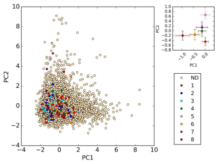

cacr

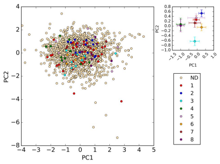

caje

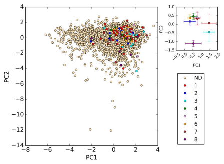

aqae

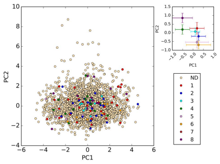

aqae

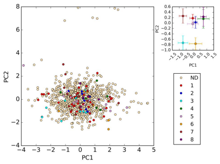

chtr

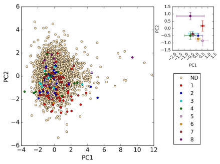

clac

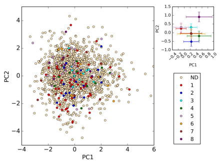

frno

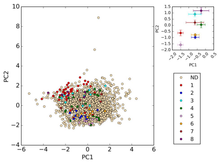

funu

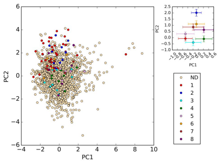

aqae

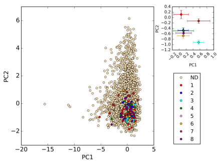

aqae

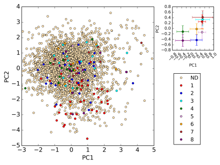

funu

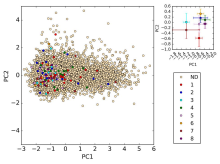

melo

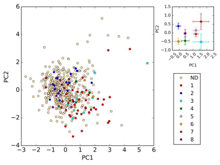

myge

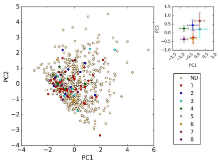

mypn

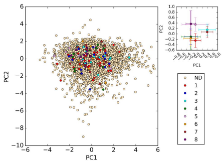

mytu

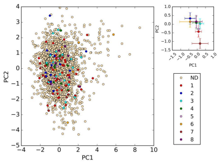

pogi

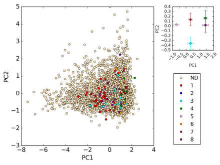

raso

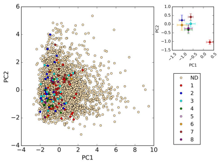

spwi

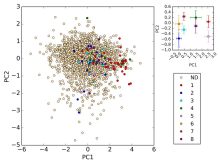

stau

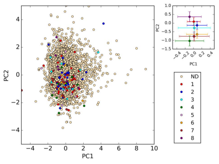

sysp

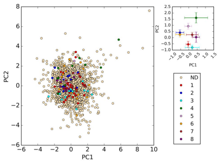

thma

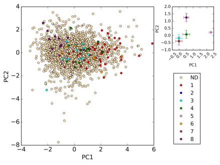

vich

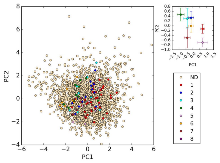

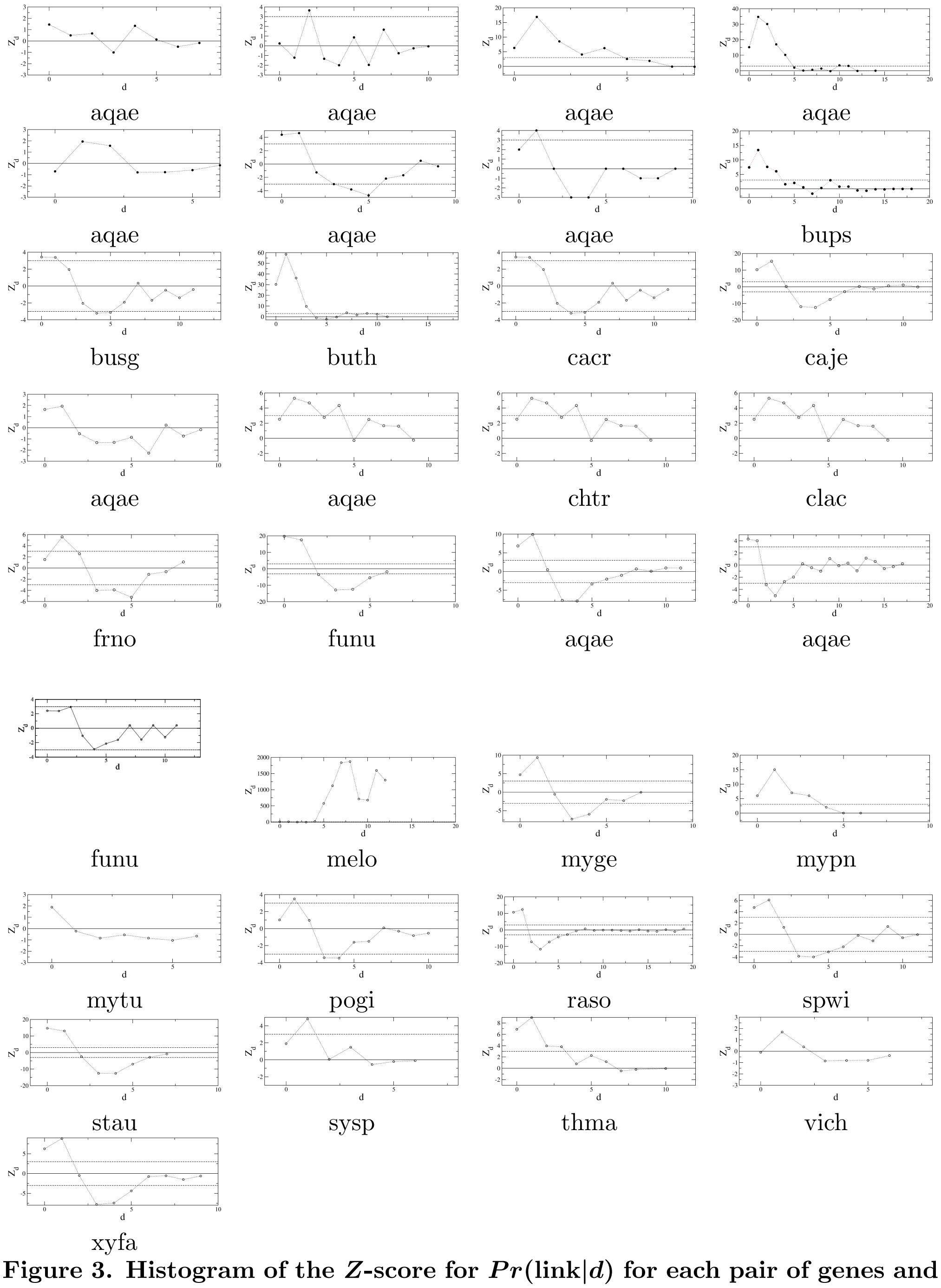


**Figure 3. Histogram of the $Z$-score for $Pr(\text{link}|d)$ for each pair of genes and their respectively encoded proteins.** $d$ is the Euclidean distance between pairs of genes in the space of the first two PCA components of codon bias, and $Pr(\text{link}|d)$ is the conditional probability of having a link in the PIN between two proteins given that their encoding genes are localized within a distances $d$ in the PC1-PC2 plane. The $Z$-score is obtained as $Z[Pr(\text{link}|d)] = [Pr(\text{link}|d) - \langle Pr(\text{link}|d)\rangle_\Omega]/\sigma_\Omega[Pr(\text{link}|d)]$. The gray dashed lines mark the significance interval of $\pm 3\sigma$.

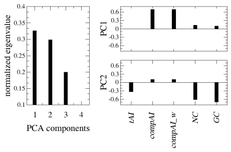 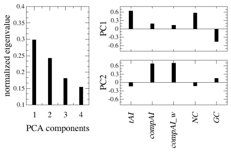 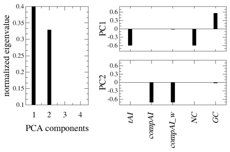 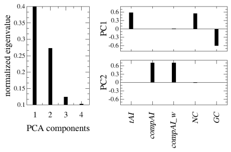

aqae aqae aqae aqae

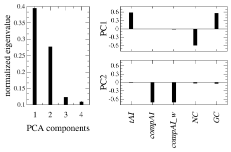 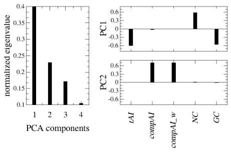 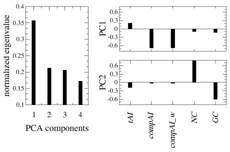 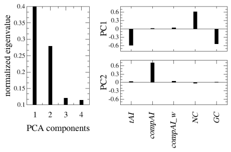

aqae aqae aqae bups

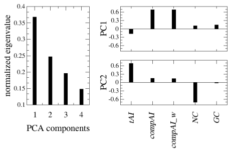 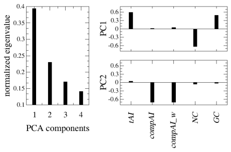 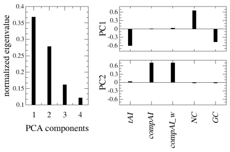 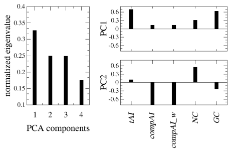

busg buth cacr caje

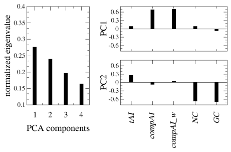 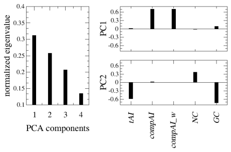 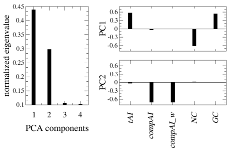 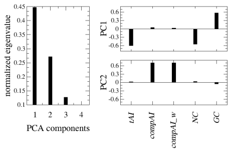

chtr clac aqae aqae

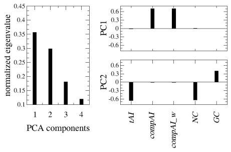 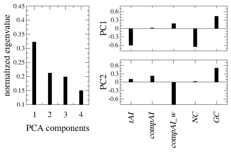 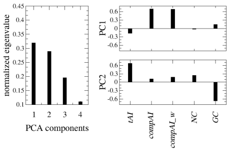 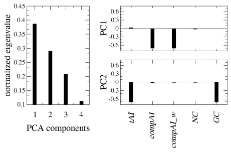

frno funu aqae aqae

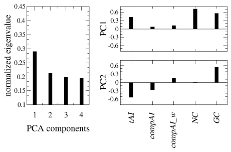 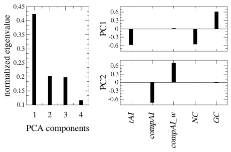 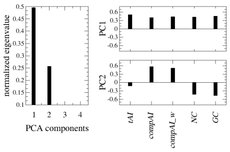 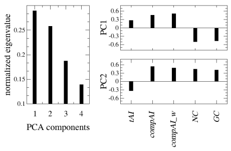

melo melo myge mypn

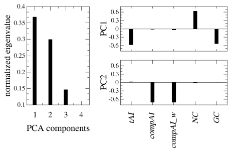 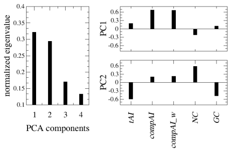 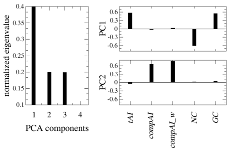 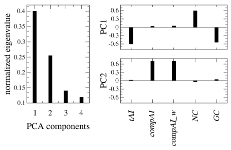

mytu pogi raso spwi

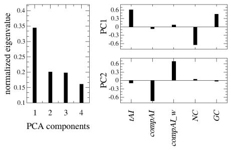 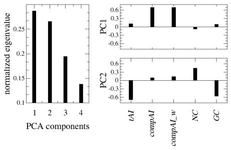 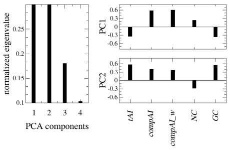 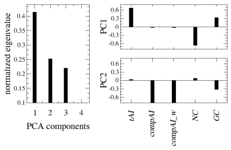

stau sysp thma vich

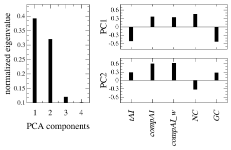
